# Plant breeders should be determining economic weights for a selection index instead of using independent culling for choosing parents in breeding programs with genomic selection

**DOI:** 10.1101/500652

**Authors:** Lorena G. Batista, R. Chris Gaynor, Gabriel R. A. Margarido, Tim Byrne, Peter Amer, Gregor Gorjanc, John M. Hickey

## Abstract

In the context of genomic selection, we evaluated and compared recurrent selection breeding programs using either index selection or independent culling for selection of parents. We simulated a clonally propagated crop breeding program for 20 cycles of selection using either independent culling or an economic selection index with two unfavourably correlated traits under selection. Cycle time from crossing to selection of parents was kept the same for both strategies. Our results demonstrate that accurate knowledge of the economic importance of traits is essential even when performing independent culling. This is because independent culling achieved its optimum genetic gain when the culling threshold for each trait varied accordingly to the economic importance of the traits. When gains from independent culling were maximised, the efficiency of converting genetic diversity into genetic gain of both selection methods were equivalent. When the same proportion selected of 10% for each trait was used instead of optimal culling levels, index selection was 10%, 128% and 310% more efficient than independent culling when T2 had a relative economic importance of 1.0, 2.5 and 5.0, respectively. Given the complexity of estimating optimal culling levels and the fact that the gains achieved with independent culling are, at most, equivalent to index selection, the use of an economic selection index is recommended for multi-trait genomic selection.

## Introduction

Crop breeding seeks to develop improved cultivars. Besides high yield levels, a successful cultivar in many crops must meet minimal standards for several other traits that are economically important, such as pest and disease resistance and product quality. Traits are often unfavourably correlated with each other (e.g., Kwon and Torrie 1964; Meredith and Bridge 1971; Erskine et al. 1985; Kato and Takeda 1996; Triboi et al. 2006). When traits are antagonistically correlated, selection for one trait causes an undesired economic response in the other trait (Falconer et al. 1996; Bernardo 2010). This makes breeding to simultaneously improve multiple traits complicated.

Independent culling and the use of a selection index are two commonly used methods in plant breeding programs for selecting on multiple traits (Bernardo 2010). Independent culling involves establishing minimum standards (i.e., culling levels) for each trait and only selecting individuals that meet these minimum standards. The thresholds can be set according to a specific selection intensity or a specific value, such as a value relative to an agronomic check. The application of independent culling can be on multiple traits simultaneously or on individual traits sequentially. The selection index method involves selection for all traits simultaneously based on a linear or non-linear combination of individual traits weighted by their importance for the breeding objective (Hazel and Lush 1942).

Theoretically, the selection index is the most efficient method of selection for multiple traits (Hazel and Lush 1942; Young 1961). However, independent culling can achieve nearly equivalent efficiencies using optimised thresholds (Xu and Muir 1991). Independent culling is less efficient, because when strictly applied it will not select individuals below the threshold for only one trait despite being exceptional for all other traits, while the use of a selection index makes it possible to retain those individuals (Bernardo 2010).

When cost is considered, independent culling can be more efficient than a selection index (Xu and Muir 1991). This is because independent culling does not require phenotypes for all individuals and traits at one time, whereas strict application of a selection index requires phenotypes for all traits. This benefit is particularly valuable to plant breeders, because early stages of the breeding program often have a very large number of individuals. Phenotyping all individuals for all traits is likely to be logistically and financially infeasible. For example, some traits have a high measurement cost, such as bread quality in wheat, so that they cannot be measured on a large number of individuals. Further, some traits can only be measured on older plants, such as lifetime production in sugarcane, or on a plot or group basis. Delaying selection until these traits become available would be effectively equivalent to random selection, because the breeder would have to reduce the overall size of the early stage. Thus, practical constraints require at least some use of independent culling on traits that can be phenotyped simply/quickly and at a lower cost in breeding programs utilising phenotypic selection.

The use of genomic selection in plant breeding may render the cost efficiency benefit of independent culling obsolete if all early generation individuals are genotyped. This is because genomic selection allows for accurate prediction of all traits at once (Meuwissen et al. 2001). While genotyping all early generation individuals is not standard in most current breeding programs, it may become so in the future. This is likely to be the case if breeding programs adopt a two-part strategy to breeding that explicitly splits breeding programs into a rapid cycling, genomic selection guided, population improvement part tasked with developing new germplasm and a product development part focused on developing new varieties. Simulations of these breeding programs suggest they can deliver considerably more genetic gain than more conventional breeding programs (Gaynor et al. 2017).

Several studies have already discussed the benefits of incorporating genomic selection strategies into crop breeding programs (Bernardo and Yu 2007; Heffner et al. 2009; Gaynor et al. 2017; Hickey et al. 2017). However, to our knowledge, there have been no studies to date that have investigated the gains over several generations in a recurrent selection breeding program using either a selection index or independent culling, at least in the context of genomic selection. We used simulations of recurrent breeding programs to evaluate and compare both strategies for 20 cycles of selection. The purpose of these simulations was to quantify the magnitude of the difference between optimally set independent culling levels and an optimal selection index. The simulations also investigated the sensitivity of independent culling to sub-optimal culling levels.

## Material and Methods

Stochastic simulations of entire breeding programs for multiple traits were used to compare the genetic gains in a breeding program using independent culling levels and a breeding program using an economic selection index for selection of parents. In the independent culling approach, selection was performed for one trait at a time at each stage of selection. A clonally propagated crop species was considered. In breeding programs for clonally propagated species, all the genotypes in the F_1_ population are candidate clones to be released as cultivars or used as parents in the next breeding cycle (Grüneberg et al. 2009). The methods were compared using the average of fifty replicates, each replicate consisting of: i) a burn-in phase shared by both strategies so that each strategy had an identical, realistic starting point; and ii) an evaluation phase that simulated future breeding with different breeding strategies. The burn-in phase consisted of 20 years of breeding using independent culling for the selection of parents and the evaluation phase consisted of 20 cycles of selection using either independent culling or index selection.

## Genome sequence

For each replicate, a genome consisting of 10 chromosome pairs was simulated for the hypothetical clonally propagated plant species. These chromosomes were assigned a genetic length of 1.43 Morgans and a physical length of 8×10^8^ base pairs. Sequences for each chromosome were generated using the Markovian Coalescent Simulator (Chen et al. 2009) and AlphaSimR (Gaynor et al.). Recombination rate was inferred from genome size (i.e. 1.43 Morgans / 8×10^8^ base pairs = 1.8×10^−9^ per base pair), and mutation rate was set to 2×10^−9^ per base pair. Effective population size was set to 50, with linear piecewise increases to 1,000 at 100 generations ago, 6,000 at 1,000 generations ago, 12,000 at 10,000 generations ago, and 32,000 at 100,000 generations ago.

## Founder genotypes

Simulated genome sequences were used to produce 50 founder genotypes. These founder genotypes served as the initial parents in the burn-in phase. This was accomplished by randomly sampling gametes from the simulated genome to assign as sequences for the founders. Sites that were segregating in the founders’ sequences were randomly selected to serve as 1,000 causal loci per chromosome (10,000 across the genome in total). To simulate genetic correlations between traits, the traits were treated as pleiotropic and the additive effects of the causal loci alleles were sampled from a multivariate normal distribution with mean 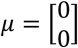 and desired values of correlation.

## Estimated breeding values

The true genetic value of the simulated traits was determined by the summing of its causal loci allele effects. The matrix **E** with the estimated breeding values of the traits for each individual in the population was obtained according to the formula:

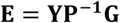

Where **Y** is the matrix of phenotypes simulated by adding random error to the true genetic values of the traits, where rows correspond to individuals in the population and columns correspond to traits. The random error was sampled from a multivariate normal distribution with mean 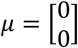 and zero covariance, with variance values tuned to achieve a target level of accuracy for both traits. **P** is the phenotypic variance-covariance matrix of the traits, and **G** is the genetic variance-covariance matrix for the traits.

## Breeding methods

The simulations modelled breeding for two component traits (T1 and T2) that were improved using either independent culling or an economic selection index. With both strategies, an F_1_ population of 5,000 individuals was generated by randomly crossing the individuals in the crossing block (Parents). With independent culling, selection was carried out in two stages: a proportion of individuals was selected first based on T1 and then, from this proportion, the parents of the next breeding cycle were selected based on T2. With the selection index approach, the F_1_ individuals with the highest values for the index trait were selected as parents of the next breeding cycle. The index trait was the sum of the estimated breeding values for each trait weighted by their economic importance. The number of selected parents (50 parents) and the cycle time from crossing to selection of new parents was kept the same for both strategies, so the comparisons between them reflect only the differences due to the method of selection. For simulation of breeding programs, we used the R package AlphaSimR (Gaynor et al.).

## Simulated scenarios

The selection index and independent culling methods were compared in a set of scenarios that aimed to assess the relative performance of the methods under different levels of accuracy of selection, and relative economic importance of T2. A summary of all simulated scenarios is shown in Table 1.

**Table 1.** Summary of parameters simulated in all comparison scenarios of recurrent selection breeding programs using either independent culling or selection index with two traits

| Scenario | Selected Proportion |  | Genetic correlation | Relative economic importance of Trait 2 | Accuracy |
| --- | --- | --- | --- | --- | --- |
|  | Trait 1 | Trait 2 |  |  |  |
| 1 | Optimum | Optimum | -0.5 | 1.0 | 0.3 |
| 2 | Optimum | Optimum | -0.5 | 1.0 | 0.5 |
| 3 | Optimum | Optimum | -0.5 | 1.0 | 0.9 |
| 4 | Optimum | Optimum | -0.5 | 1.0 | 0.7 |
| 5 | Optimum | Optimum | -0.5 | 2.5 | 0.7 |
| 6 | Optimum | Optimum | -0.5 | 5.0 | 0.7 |
| 7 | 10% | 10% | -0.5 | 1.0 | 0.7 |
| 8 | 10% | 10% | -0.5 | 2.5 | 0.7 |
| 9 | 10% | 10% | -0.5 | 5.0 | 0.7 |

For one set of scenarios we simulated four levels of accuracy (0.3, 0.5, 0.7, and 0.99), assigned the same economic importance for both traits, and an unfavourable initial genetic correlation of −0.5 between traits. In another set of scenarios, we varied the relative economic importance of T2, but fixed selection accuracy to 0.7 and set an unfavourable initial genetic correlation of −0.5 between traits. Here, three levels of relative economic importance were simulated. T1 was given an economic importance of 1.0 and T2 an economic importance of either 1.0, 2.5 or 5.0. For each level of relative economic importance, we simulated: i) scenarios where the proportion selected was the same (10%) when selecting for both traits, and ii) scenarios where the proportions selected were set to achieve optimal culling levels (i.e., optimal independent culling). To achieve optimal culling levels, in each cycle of selection we chose the proportion selected for each trait that maximised the genetic gain. To find the optimal proportions, we fixed the number of parents selected (50 parents) and found the number of individuals to be selected in the first culling stage that maximized parents’ economic value (i.e., index trait).

## Comparison

The comparisons were made in terms of: i) genetic gain ii) genetic diversity, iii) the efficiency of converting genetic diversity into genetic gain for the index; and iv) genetic correlation between traits. For genetic gain and genetic diversity, we report values based on the individuals in the crossing block (parents) at each cycle of selection. We measured genetic gain as the increment in genetic mean (average of true genetic values) compared to the genetic mean in year 20. We measured genetic diversity with genetic standard deviation and genic standard deviation. We calculated genetic standard deviation as standard deviation of true genetic values. We calculated genic standard deviation as 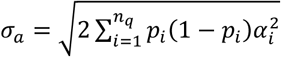, where *n_q_* is the number of causal loci and *p_i_* and *α_i_* are, respectively, allele frequency and allele substitution effect at the *i*-th causal locus.

To measure efficiency, genetic mean and genic standard deviation were standardized to mean zero and unit standard deviation in year 20. We measured efficiency of converting genetic diversity into genetic gain by regressing the achieved genetic mean 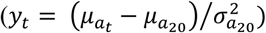 on lost genetic diversity 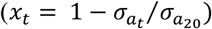, i.e., *y_t_ = α + bx_t_ + e_t_*, where *b* is efficiency (Gorjanc et al. 2017). We estimated efficiency with robust regression using function rlm() in R (Venables and Ripley 2002).

For genetic correlation, we report the correlation between the true genetic values of T1 and T2. We calculated this metric on the individuals in the F_1_ population at each cycle of selection.

## Results

Overall the results show that index selection provided consistent genetic gains and was equivalent to independent culling in terms of genetic gains and efficiency when optimal culling levels were used. Index selection performed better than independent culling in scenarios where independent culling levels were suboptimal.

We have structured the description of the results in two parts, corresponding to how the relative performance of the selection methods was affected by: i) the accuracy of selection, and ii) the relative economic importance of traits.

## Accuracy of selection

The results show that increases in accuracy accentuated the differences in the genotypes being selected by either independent culling or index selection. This is shown in Fig. 1, where the genotypes selected as parents by each selection method are highlighted. Lower levels of accuracy led to a more diffuse cluster of selected genotypes and, with increasing selection accuracy, the cluster of selected genotypes approached what was expected for each method of selection (Bernardo, 2010).

**Fig 1.**
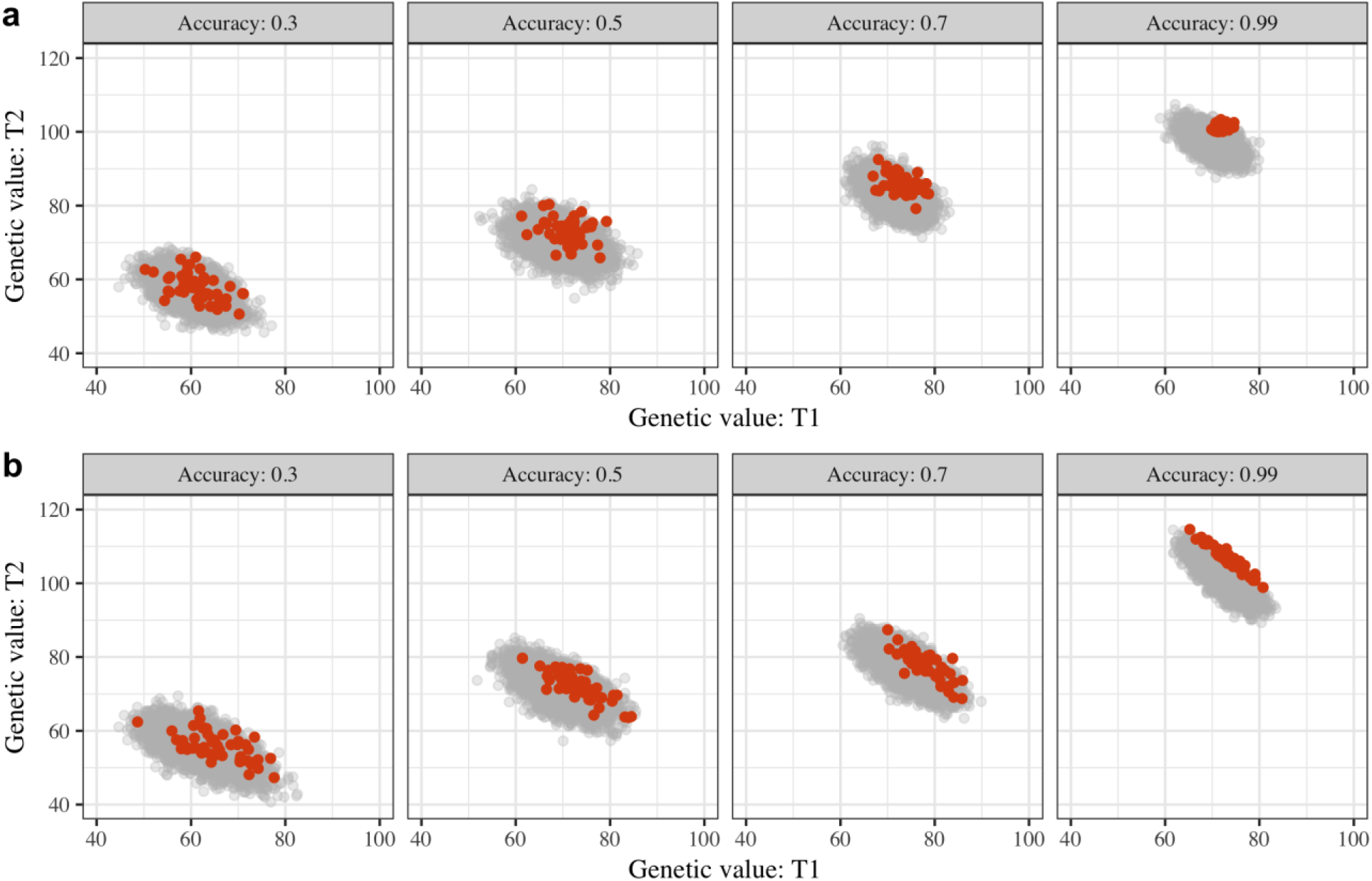
Scatterplots of true genetic values for Trait 1 (T1) and Trait 2 (T2) of the genotypes in the F_1_ population (grey) and genotypes selected as parents (orange) in the third cycle of selection using either independent culling (**a**) or a selection index (**b**) with different levels of accuracy

Fig. 2 shows the change in the genetic correlation between the component traits for both independent culling and index selection over 20 cycles of selection at different levels of accuracy. Both selection methods resulted in the correlation between traits becoming increasingly unfavourable over the cycles of selection. For both methods, the change in the genetic correlation was higher with higher values of accuracy. Compared to independent culling, index selection led to larger changes in the genetic correlation between the two traits. After 20 cycles of selection with accuracy of 0.3, independent culling led to a genetic correlation that was 9% more unfavourable compared to the genetic correlation in cycle 0, while index selection led to a genetic correlation that was 17% more unfavourable compared to the genetic correlation in cycle 0. After 20 cycles of selection with accuracy of 0.99, independent culling led to a genetic correlation that was 29% more unfavourable compared to the genetic correlation in cycle 0, while index selection led to a genetic correlation that was 64% more unfavourable compared to the genetic correlation in cycle 0.

**Fig 2.**
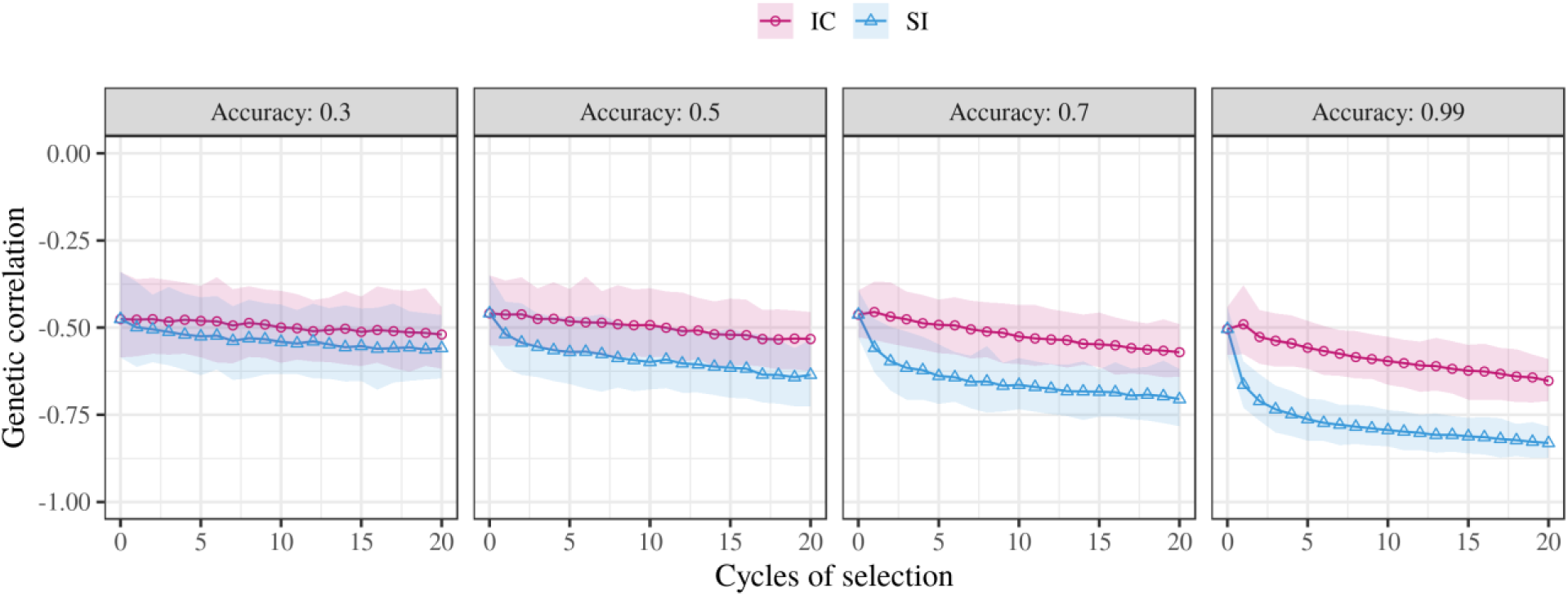
Change in genetic correlation (mean and 95% confidence interval) between traits in the F_1_ population over 20 cycles of selection using either optimal independent culling (IC) or a selection index (SI) with different levels of accuracy, and Trait 2 relative economic importance of 1.0

The change of genetic mean in parents for the component traits and the index trait over the cycles of selection using each method is shown in Fig. 3. For both methods, the genetic gains for the component traits and the index trait increased with higher values of accuracy. In general, the selection index method and independent culling with optimal culling levels led to equivalent genetic gains for the component traits and the index trait. Only in the scenario with 0.99 accuracy did index selection lead to a slightly higher genetic gain compared to that achieved with optimal independent culling. For the index trait, after 20 cycles of selection with accuracy of 0.99, index selection had a genetic gain 4% higher than the genetic gain achieved with independent culling.

**Fig 3.**
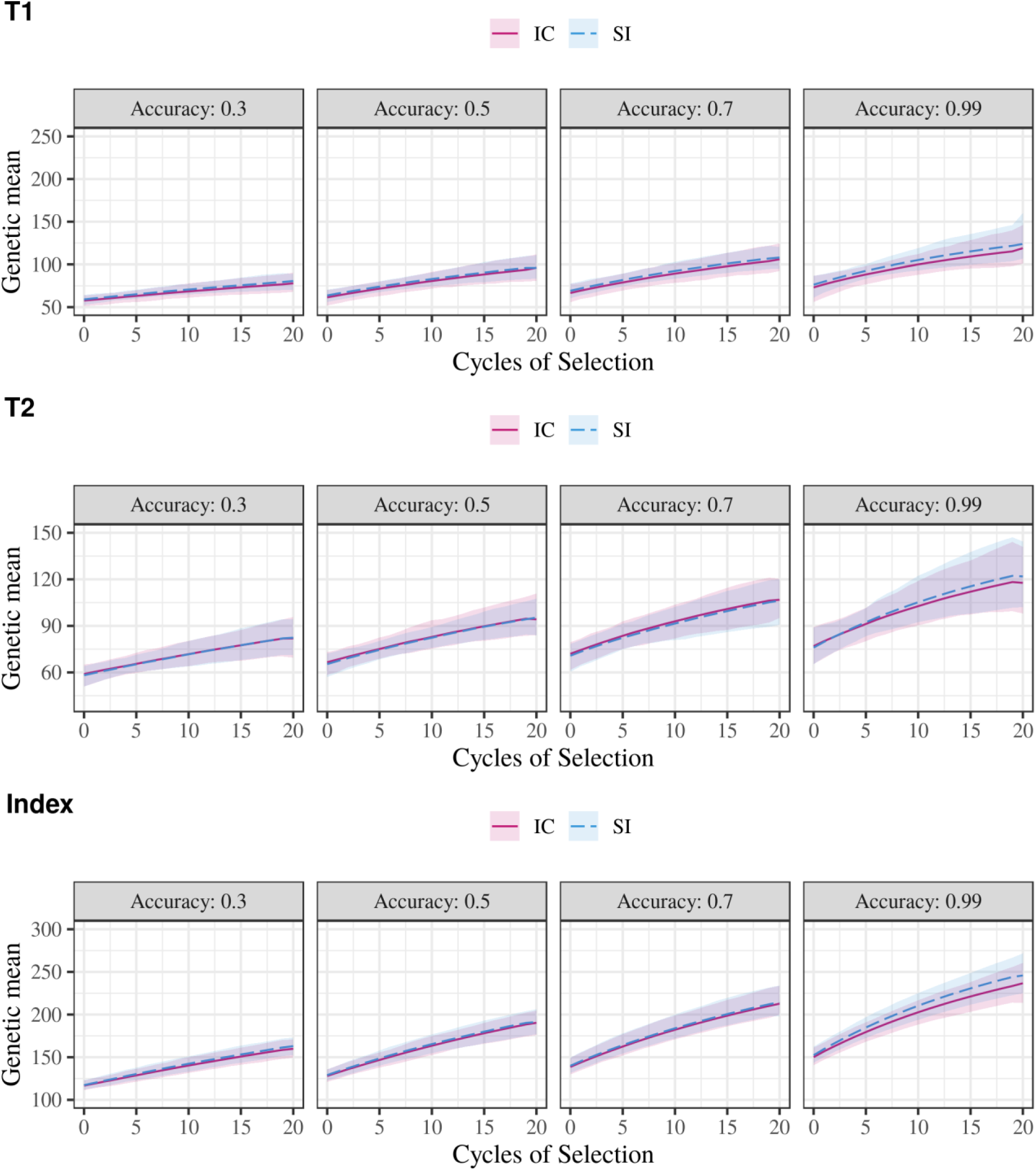
Change in genetic mean for Trait 1 (T1), Trait 2 (T2) and Index Trait (Index) over 20 cycles of selection using either optimal independent culling (IC) or a selection index (SI) with different levels of accuracy, unfavourably correlated traits, and T2 relative economic importance of 1.0

Table 2 shows the genetic standard deviation of parents in cycle 20 and the loss in genetic standard deviation in cycle 20 compared to the genetic standard deviation in cycle 0 for the component traits and the index trait. The change of genetic diversity in parents for the component traits and the index trait over the cycles of selection using each method is shown in Supplementary material 1 (Fig S1.1). For the component traits, when using index selection, the genetic standard deviation showed an initial increase in the first few cycles of selection followed by a gradual decrease in the subsequent cycles. When using independent culling, the decrease in the genetic standard deviation of the component traits was continual over the cycles of selection. Both of these trends were more obvious with increasing values of accuracy. For all values of accuracy, independent culling led to a higher loss in the genetic standard deviation of the component traits compared to the index selection. For T1 and T2, independent culling with accuracy of 0.3 led to a loss of genetic standard deviation that was 6% and 5% higher than the loss of genetic standard deviation observed for index selection, respectively. With accuracy of 0.99, for T1 and T2 independent culling led to a loss of genetic standard deviation that was 65% and 51% higher than the loss of genetic standard deviation observed for index selection, respectively. For the index trait, both methods led to equivalent values of genetic standard deviation. With accuracies of 0.3 and 0.99, index selection led to a loss in the genetic standard deviation of the index trait that was 3% higher compared to the loss of genetic standard deviation observed using independent culling, respectively.

**Table 2.** Mean genetic standard deviation (Genetic SD) of parents in cycle 20 and loss in genetic standard deviation in cycle 20 in comparison to the genetic standard deviation in cycle 0 (Loss over cycle 0) for trait 1 (T1), trait 2 (T2) and the index trait using either optimal independent culling or index selection with different levels of accuracy, unfavourably correlated traits, and T2 relative economic importance of 1.0

| Accuracy | Independent culling |  |  |  |  |  |
| --- | --- | --- | --- | --- | --- | --- |
|  | T1 |  | T2 |  | Index trait |  |
|  | Genetic SD (cycle 20) | Loss over cycle 0 | Genetic SD (cycle 20) | Loss over cycle 0 | Genetic SD (cycle 20) | Loss over cycle 0 |
| 0.3 | 3.51 (0.08)* | -17% | 3.68 (0.08) | -16% | 3.57 (0.06) | -22% |
| 0.5 | 2.56 (0.06) | -30% | 2.45 (0.04) | -28% | 2.69 (0.05) | -32% |
| 0.7 | 1.65 (0.04) | -42% | 1.64 (0.03) | -37% | 1.88 (0.04) | -45% |
| 0.99 | 0.45 (0.01) | -68% | 0.45 (0.01) | -55% | 0.74 (0.02) | -62% |

| Accuracy | Index Selection |  |  |  |  |  |
| --- | --- | --- | --- | --- | --- | --- |
|  | T1 |  | T2 |  | Index trait |  |
|  | Genetic SD (cycle 20) | Loss over cycle 0 | Genetic SD (cycle 20) | Loss over cycle 0 | Genetic SD (cycle 20) | Loss over cycle 0 |
| 0.3 | 3.80 (0.09) | -11% | 4.00 (0.09) | -11% | 3.66 (0.08) | -19% |
| 0.5 | 3.19 (0.08) | -17% | 3.19 (0.07) | -14% | 2.57 (0.06) | -33% |
| 0.7 | 2.69 (0.06) | -16% | 2.60 (0.06) | -18% | 1.86 (0.04) | -41% |
| 0.99 | 1.93 (0.4) | -3% | 1.91 (0.04) | -4% | 0.51 (0.01) | -59% |

\* standard errors of the estimates are presented in parenthesis

Table 3 shows the genic standard deviation of parents in cycle 20 and the loss in genic standard deviation in cycle 20 compared to the genic standard deviation in cycle 0 for the component traits and the index trait. The values of genic standard deviation of T1, T2, and the index trait were equivalent. The highest difference between methods in the loss in genic standard deviation was 1% for all values of accuracy, except with accuracy of 0.99. With 0.99 accuracy, for T1, T2 and the index trait, index selection led to a loss in the genic standard deviation that was 3% higher compared to the loss of genic standard deviation observed using independent culling.

**Table 3.** Genic standard deviation (Genic SD) of parents in cycle 20 and loss in genic standard deviation in cycle 20 in comparison to the genic standard deviation in cycle 0 (Loss over cycle 0) for trait 1 (T1), trait 2 (T2) and the index trait using either optimal independent culling or index selection with different levels of accuracy, unfavourably correlated traits, and T2 relative economic importance of 1.0

| Accuracy | Independent culling |  |  |  |  |  |
| --- | --- | --- | --- | --- | --- | --- |
|  | T1 |  | T2 |  | Index trait |  |
|  | Genic SD (cycle 20) | Loss over cycle 0 | Genic SD (cycle 20) | Loss over cycle 0 | Genic SD (cycle 20) | Loss over cycle 0 |
| 0.3 | 3.94 (0.06)* | -15% | 4.11 (0.07) | -15% | 4.04 (0.05) | -16% |
| 0.5 | 3.48 (0.06) | -24% | 3.41 (0.05) | -24% | 3.44 (0.04) | -25% |
| 0.7 | 2.94 (0.04) | -34% | 2.89 (0.04) | -34% | 2.89 (0.04) | -34% |
| 0.99 | 2.35 (0.04) | -42% | 2.35 (0.04) | -42% | 2.33 (0.04) | -43% |

| Accuracy | Index Selection |  |  |  |  |  |
| --- | --- | --- | --- | --- | --- | --- |
|  | T1 |  | T2 |  | Index trait |  |
|  | Genic SD (cycle 20) | Loss over cycle 0 | Genic SD (cycle 20) | Loss over cycle 0 | Genic SD (cycle 20) | Loss over cycle 0 |
| 0.3 | 3.92 (0.06) | -16% | 4.08 (0.07) | -16% | 4.02 (0.05) | -16% |
| 0.5 | 3.44 (0.06) | -25% | 3.37 (0.05) | -25% | 3.39 (0.05) | -26% |
| 0.7 | 2.92 (0.05) | -34% | 2.88 (0.05) | -34% | 2.87 (0.04) | -35% |
| 0.99 | 2.21 (0.04) | -45% | 2.22 (0.04) | -45% | 2.17 (0.03) | -46% |

\* standard errors of the estimates are presented in parenthesis

## Relative economic importance of traits

Fig. 4 shows the efficiency of converting genetic diversity into genetic gain for the index trait when the relative economic importance of T2 varies. Independent culling was compared to index selection using either optimal culling levels or selection with the same proportion of plants selected (10%) for each trait. Index selection had the highest efficiency and most gain for all levels of economic importance. The efficiency and gain for optimal independent culling levels was nearly equivalent to index selection. The efficiency and gain for selecting the same proportion of plants for both traits was worse than index selection for all levels of relative economic importance. Index selection was 10%, 128% and 310% more efficient than independent culling using the same proportion of selected plants for relative economic importance of 1.0, 2.5 and 5.0, respectively.

**Fig 4.**
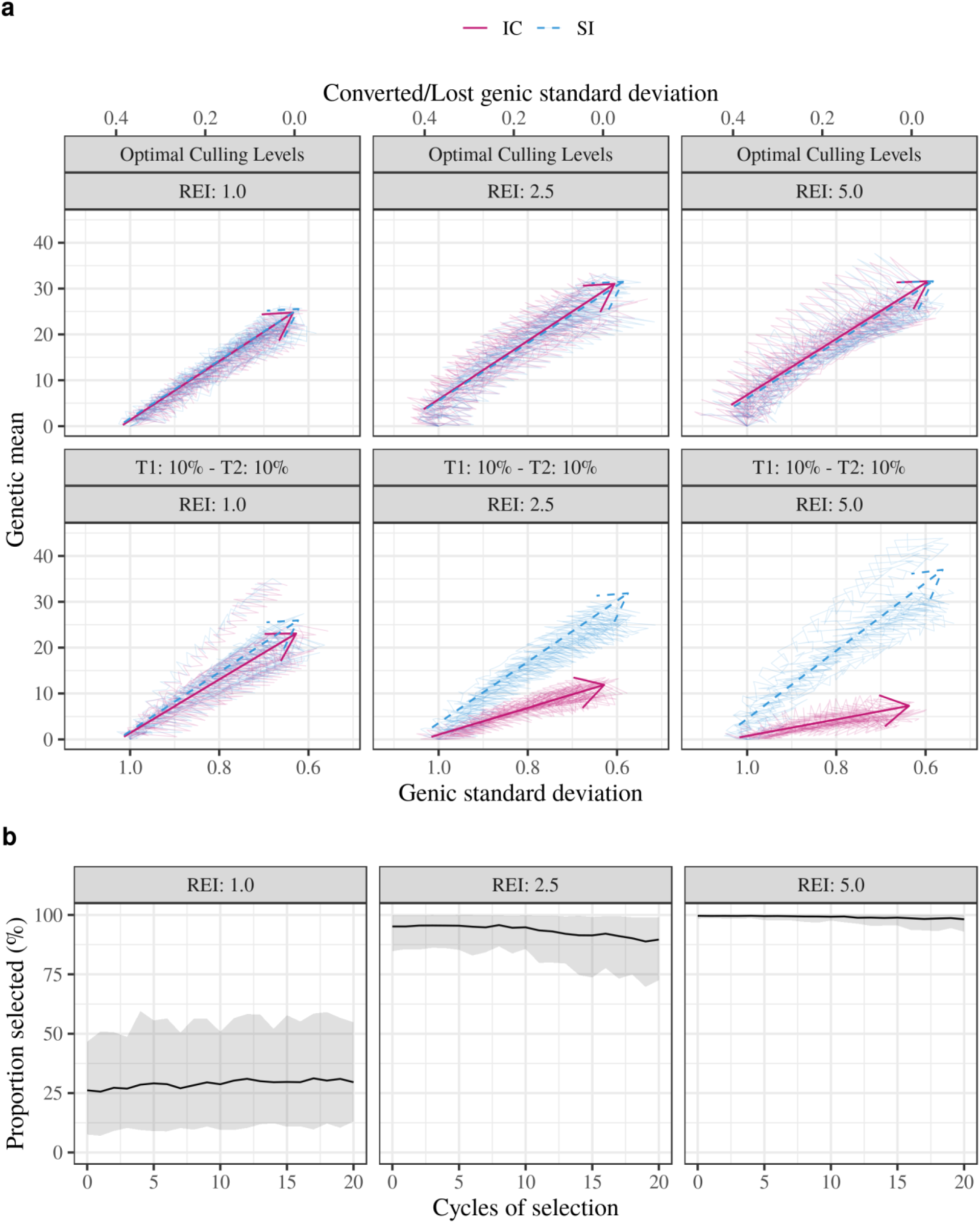
Change of genetic mean and genic standard deviation for the index trait across 20 cycles of selection using either independent culling (IC) or a selection index (SI) under three levels of relative economic importance (REI) and using either the same proportion selected (10%) for Trait 1 (T1) and Trait 2 (T2) or optimal culling levels for each level of relative economic importance of T2 (a); and proportion selected (mean and 95% confidence interval) for T1 used to achieve optimal culling levels over the 20 cycles of selection (b). Traits are unfavourably correlated (−0.5). Individual replicates are shown by thin lines and a mean regression with a time-trend arrow. Values of genetic mean and genic standard deviation shown are standardized to mean zero and unit standard deviation in cycle 0

Fig. 4 also shows the proportion of plant selected for T1 under optimal independent culling over the different levels of economic importance for T2. The mean proportion selected for T1 only varied slightly over the cycles of selection. The means were 29%, 93%, and 99% for relative economic importance of 1.0, 2.5, and 5.0, respectively. The variation about those means was largest with relative economic importance of 1.0 and smallest with relative economic importance of 5.0.

## Discussion

This study evaluated and compared recurrent selection breeding programs that either use index selection or independent culling for the selection of parents by genomic selection. Overall the results show that using index selection is either better or equivalent to independent culling in this context. Index selection outperformed independent culling when sub-optimal culling levels were used. Our results demonstrate that accurately assessing the economic importance of the traits is essential regardless of the method of selection being used.

The main difference between index selection and independent culling is that, when using index selection, genotypes that are exceptional for one of the traits under selection are more likely to be selected even though their performance for other traits is average. This can be seen in Fig. 1, with the cluster of individuals selected as parents with the index method including individuals that are more contrasting for the two traits under selection compared to the individuals selected with independent culling. The main implications of this are in the way each method affects the correlation between traits and the genetic diversity over cycles of recurrent selection. We discuss each of these aspects in the following two sections. In the third section, we discuss how the relative economic importance of the traits can affect the relative performance of the methods. Lastly, we discuss the implications of our results for modern plant breeding programs which deploy genomic selection.

## Methods of selection and genetic correlation between traits

The results show that, after only a few cycles of selection, index selection generates F_1_ populations with a more unfavourable genetic correlation between traits than the F_1_ populations generated by independent culling (Fig 2). An explanation for the faster decrease of the genetic correlation observed with index selection is that the index is a linear combination of component traits. As shown by Bulmer (1971), selection on a linear combination leads to negative covariances between components (i.e., Bulmer effect). Consequently, the same principle applies to the component traits and index selection, with index selection leading to an unfavourable genetic correlation between the component traits (Tallis 1987; Itoh 1991).

In general, genetic gains in multi-trait selection, regardless of the method of selection, are expected to be higher when the correlation between traits is favourable and lower when this correlation is unfavourable (Young 1961). As index selection generated F_1_ populations with more unfavourable genetic correlation between traits than independent culling, the genetic gains for index selection were potentially lower than for independent culling. Nevertheless, despite index selection being carried out under increasingly unfavourable genetic correlations over the cycles, the genetic gains obtained for the index trait were equivalent to the gains obtained using independent culling (Fig. 3).

Unfavourable genetic correlations are the most challenging scenario for breeders. When traits are unfavourably correlated, selection on one trait results in response in an undesired direction for the other trait. When these correlations are due to pleiotropy, they cannot be broken with repeated cycles of recombination. This case is likely pervasive in several crops, e.g., grain yield and protein content in cereal crops (Duvick and Cassman 1999; Rharrabti et al. 2001; Rotundo et al. 2009), quality and disease resistance in forage crops (Casler and Vogel 1999), and yield and disease resistance in barley (Smedegaard-Petersen and Tolstrup 1985). However, the extent of genetic correlation and pleiotropy in these examples is unknown because unfavourable genetic correlations between the traits could also be, at least partly, induced by selection, as demonstrated in this study.

## Methods of selection and genetic diversity over cycles of selection

According to Bulmer (1971), reduction in the genetic variance due to selection stems mostly from the build-up of negative linkage disequilibrium between causal loci when selection is performed. This can be seen by comparing genetic and genic variation (Table 2 and Table 3, respectively). Genic variation is a function of the allele frequencies and the allele substitution effect only, and thus is not affected by changes in linkage disequilibrium. The results in Table 3 show that the loss of genic standard deviation of the component traits and index trait are not greatly affected by the method of selection. Also, the method of selection did not greatly affect the trait means, as shown in Fig. 3. This indicates that, in terms of allele frequencies, there was little difference in the parents selected by either independent culling or the selection index method in situations similar to our simulation. Therefore, the difference between the selection methods derives from how they induce and exploit linkage disequilibrium between the causal variants of the component traits. Specifically, as shown in Table 2, independent culling induced a greater degree of negative linkage disequilibrium between the causal variants of the component traits resulting in those traits having less genetic variation. A deviation from this result is expected with more intense selection schemes and more component traits selected in successive stages, which would induce larger changes in allele frequencies due to drift. As a consequence, differences between index selection and independent culling would be accentuated. Cowling and Li (2018) simulated and compared wheat breeding programs using different selection strategies under high and low selection intensities. They observed index selection resulted in higher population coancestry over cycles of selection compared to independent culling, and the difference between methods increased in scenarios with high selection intensity. Their results indicate index selection leads to a higher loss of genic standard deviation.

Somewhat surprisingly, it is possible to make an argument for the superiority of independent culling relative to a selection index on the basis of the differences observed in linkage disequilibrium. This is because independent culling produced populations with nearly equivalent mean performance, but with more consistent performance between individuals, which is demonstrated by the lower variation observed for the component traits. This property could be beneficial from a management perspective if differences in the component traits require variations in management of individuals. Breeding for plant-architecture traits in outbreeding cultivars is a good example where this property might be valuable, as having more uniform plants in the field favours mechanical harvest. However, we believe this property is more of an academic curiosity than something that will have practical application.

For simplicity and ease of implementation, our simulations consider the same genetic architecture for both traits, with both traits being controlled by a high number (10,000) of causal loci with small additive effects. Under different circumstances, such as at least one of the traits being controlled by few causal loci with higher allele substitution effects, different results could be expected. The results for the two-locus model of Bennett and Swiger (1980) show that independent culling tends to eliminate genotypes that are homozygous for alleles with low effect for one of the traits. For one pleiotropic causal locus, when both alleles are favourable for one trait and unfavourable for the other trait, both homozygous genotypes tend to be culled, and independent culling would select the heterozygous genotypes. If heterozygous genotypes were preferred, the fixation of alleles would be slower and, therefore, the loss in genic standard deviation would be lower. Our results indicate that, for highly polygenic traits, differences between methods of selection in the loss of genetic diversity are mostly due to changes in linkage disequilibrium as opposed to distinctive changes in allele frequencies. Therefore, in terms of conserving genetic diversity there was no obvious advantage for either method. Other strategies such as optimal-cross selection should be considered in order to optimize gains while also controlling the loss of genetic diversity over cycles of selection (Clark et al. 2013; Woolliams et al. 2015; Gorjanc et al. 2017; Cowling and Li 2018).

## Economic importance of the traits

In general, when using the same selection intensity for both traits, the greater the difference in the economic importance of the traits, the better index selection will perform compared to independent culling (Fig. 4). This happens because there is a combination of selection intensities for each trait that maximizes the genetic gain when performing independent culling (Hazel and Lush 1942). Finding these selection intensities when selecting for two traits in two stages of selection is complex (Young and Weiler 1960; Namkoong 1970; Cotterill and James 1981; Smith and Quaas 1982), and becomes even more complex with increasing number of traits and stages of selection (Saxton 1989; Ducrocq and Colleau 1989; Xu and Muir 1991).

The results in Fig. 4 show that independent culling approaches its maximal gain when a higher selection intensity is used for the trait with higher economic importance and a lower selection intensity is used for the trait with lower economic importance. In fact, when one trait had 5 times the economic importance of the other trait, the optimum was achieved when almost no selection was carried out for the less important trait. These results demonstrate that accurately assessing the economic importance of the traits is essential even when independent culling is performed.

Regardless of the gains achieved with independent culling being maximised, when parents are selected based on an index, equivalent gains are achieved by simply summing the values of the traits weighted by their economic importance. Once the true economic weights of the traits are quantified, index selection is much simpler than independent culling when using these weights for optimizing the genetic gains in a plant breeding program.

## Index selection in modern plant breeding programs that use genomic selection

There is little to no evidence suggesting plant breeders use analytical techniques to determine optimal independent culling thresholds and/or constructing selection indices in most plant breeding programs. More likely, the majority of breeders rely on their intuition for setting thresholds and constructing indices. Their decisions are likely guided by the performance of agronomic checks and are prone to fluctuations between seasons and individual breeders. This model has clearly been successful, because plant breeding programs have continued to deliver genetic gain. However, it is likely sub-optimal, and a more analytical approach should be adopted in the future.

The value of a more analytical approach becomes greater as genomic selection is more widely used. The results presented in this paper show a selection index is superior to independent culling when using genomic selection. These results are further supported by earlier theoretical work (Smith 1936; Hazel and Lush 1942; Young 1961). This indicates a clear preference for implementing selection indices in plant breeding.

The focus of plant breeders should be determining the economic weights for a selection index. In this paper the economic model used to select weights was implicitly assumed to be known and linear. The reality is that true economic model may be unknown to breeders and it is likely non-linear. The presence of a non-linear model does not pose a problem, because linear economic weights can be derived for improving the economic value of germplasm (Goddard 1983). However, this still requires defining the economic model. For this reason, it is our opinion that plant breeders would benefit greatly from an increased emphasis on understanding and quantifying the economics of their species. This information would greatly aid breeders in getting the most out of genomic selection.

## Conclusions

We evaluated and compared recurrent selection breeding programs using either independent culling or index selection for parent selection. The results show that, despite selection being carried out under unfavourable genetic correlations when using the selection index instead of independent culling, equivalent or higher genetic gains were achieved with index selection in all simulated scenarios. In terms of genetic diversity, the differences between methods in the studied system were driven mostly by differences in the generation of linkage disequilibrium between causal loci induced and not differences in allele frequencies. When linkage disequilibrium was not considered, both methods were equivalent in terms of loss of genetic diversity, and the differences between methods in terms of efficiency of converting genetic diversity into genetic gains mostly reflected the differences in the genetic gains obtained with each method. To obtain higher genetic gains, accurately assessing the economic importance of the traits is essential even when independent culling is performed, as optimal culling levels should be determined in order for maximum gain to be achieved. Given that optimal culling levels are complex to estimate, once the economic importance of each trait is known, maximum genetic gains are more easily achieved with index selection. Therefore, the best choice for plant breeding programs is to select parents using an economic selection index.

## Acknowledgements

This study was supported by the Coordenação de Aperfeiçoamento de Pessoal de Nível Superior (CAPES, Computational Biology Programme, Grant No. BEX 0043/17-6). The Roslin authors acknowledge the financial support from BBSRC and KWS UK, RAGT Seeds Ltd., Elsoms Wheat Ltd and Limagrain UK for the project “GplusE: Genomic selection and Environment modelling for next generation wheat breeding” (grants BB/L022141/1 and BB/L020467/1).

