## Supplementary Figure 1 for "Plant breeders should be determining economic weights for a selection index instead of using independent culling for choosing parents in breeding programs with genomic selection"

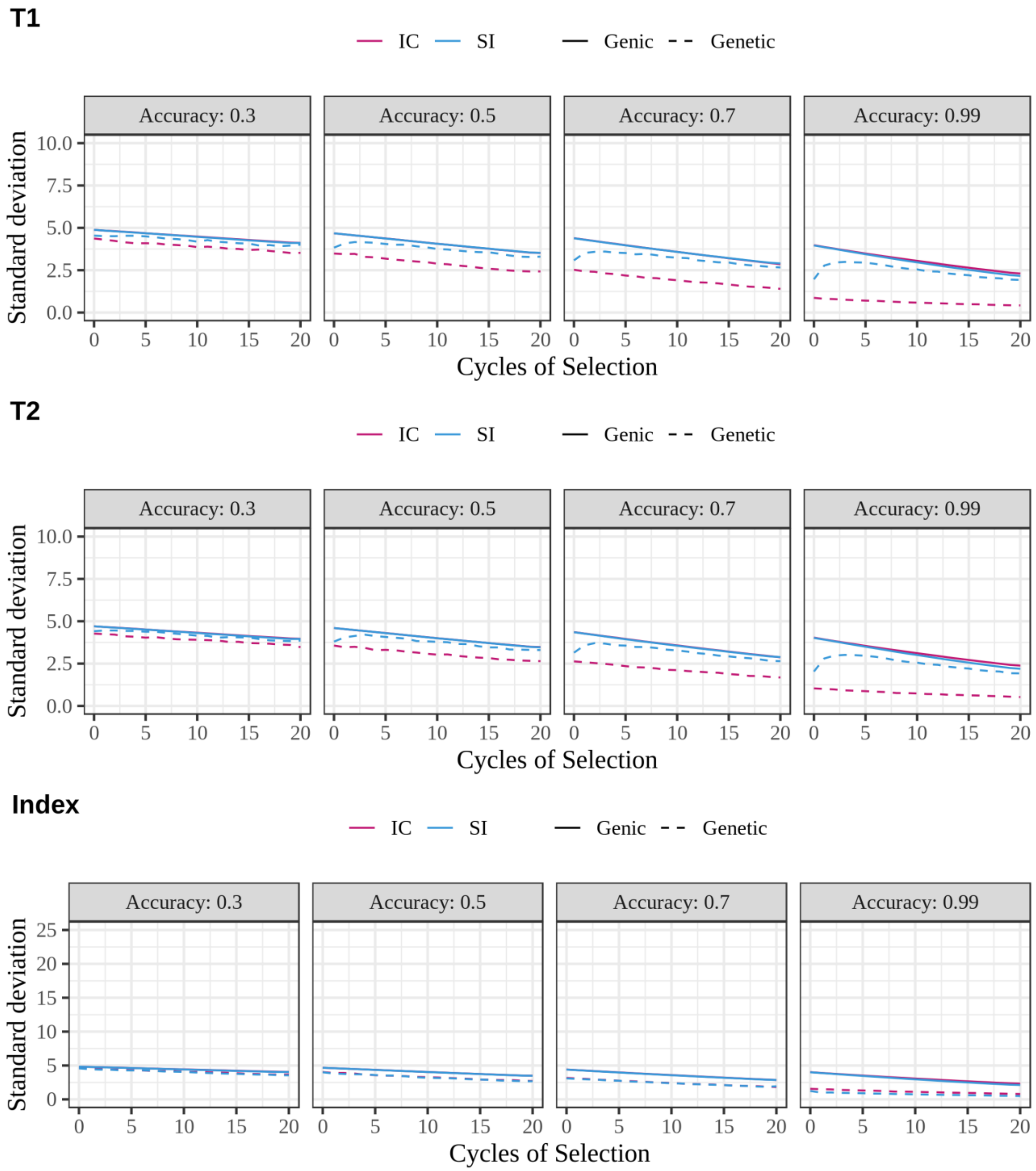


Figure S1.2 Change in genic and genetic standard deviation for Trait 1 (T1), Trait 2 (T2) and Index Trait (Index) over 20 cycles of selection using either independent culling (IC) or a selection index (SI) with different levels of accuracy, proportion selected of 10%, unfavourably correlated traits, and T2 relative economic importance of 1.0
